# Integration of Hematopoietic and Thymus-like Niches in a Human iPSC-derived Bone Marrow Organoid

**DOI:** 10.1101/2025.11.04.686649

**Authors:** Jiyoung Lee, Tomoyuki Kawasaki, Lilika Tabata, Junlong Chen, Toru Uchiyama, Satoshi Yamazaki, Akihiro Umezawa, Hidenori Akutsu

## Abstract

Human bone marrow generates blood cells but critically lacks the thymic environment required for T cell development. Developing an integrated in vitro platform that reconstitutes both functions simultaneously remains a major challenge in regenerative and immune medicine. We asked whether a synthetic marrow could be engineered to provide both capacities. Using human induced pluripotent stem cells, we created self-organizing bone marrow organoids (iBMOs) that faithfully reproduced native stromal and vascular structures and, remarkably, supported robust thymus-like T cell differentiation. iBMOs directed hematopoietic progenitors toward functional T and dendritic cells. When engrafted into immunodeficient mice, they autonomously sustained human erythropoiesis and de novo bone formation in vivo. Single-cell transcriptomics revealed a complex niche architecture, including a hybrid cluster co-expressing stem, endothelial, and stromal markers, key to understanding this dual functionality. This bioengineered organoid-stromal system unifies bone marrow and thymic functions in a single human-derived platform, offering a scalable source of immune cells for adoptive cell therapies and regenerative applications, as well as a versatile model for studying human hematopoiesis and immunity.

## Introduction

Human hematopoiesis is a hierarchically organized process through which all blood cell types are generated from a rare pool of hematopoietic stem cells (HSCs)^1^. These multipotent cells arise during embryogenesis in the aorta–gonad–mesonephros region and co-express endothelial markers such as CDH5 and SOX7 together with hematopoietic transcription factors, including RUNX1 and MYB. After birth, HSCs reside primarily in the bone marrow, where a complex network of mesenchymal stromal cells (MSCs), endothelial cells, and perivascular niches orchestrates self-renewal, lineage specification, and maturation^2–4^. MSCs contribute trophic factors, maintain tissue architecture, and regulate immune responses^5–7^. In contrast, T lymphopoiesis occurs in the thymus, where specialized stromal populations provide unique Notch-dependent cues^8^. The anatomical segregation of marrow- and thymus-dependent processes poses a major obstacle to constructing in vitro systems that reproduce the full spectrum of human hematopoiesis.

HSC transplantation remains a cornerstone therapy for hematological malignancies and immune deficiencies^9,10^, but its efficacy is limited by donor availability, HLA mismatch, and the risk of graft-versus-host disease^11–13^. Moreover, immune reconstitution is often delayed, particularly for T cells, leaving patients vulnerable to infection and relapse. These clinical challenges underscore the need for scalable, autologous platforms to generate functional hematopoietic and immune cells under physiologically relevant conditions. Organoid technology harnesses the self-organizing capacity of pluripotent stem cells to model human development in vitro^14,15^, yet no bone marrow model has recapitulated the architectural complexity of native marrow or demonstrated thymus-like T cell production. Here, we establish human induced pluripotent stem cell (iPSC)-derived bone marrow organoids (iBMOs), under extracellular matrix–free conditions. iBMOs reproduced stromal and vascular compartments, sustained hematopoietic progenitors, and displayed in vivo hematopoietic and ossifying capacity. Furthermore, we established stromal stem cell lines (iBOSS) that unexpectedly supported the differentiation of progenitors into both T cells and dendritic cells. This integrated platform unifies marrow and thymic functions in a single human-derived system, offering broad opportunities in immunology, regenerative medicine, disease modelling, and immune cell manufacturing.

## Results

### Functional iBMOs from human iPSCs

The bone marrow is a semi-solid tissue that is embedded within trabecular bone and supports hematopoiesis through a vascular microenvironment of myeloid, lymphoid, endothelial, and mesenchymal stromal cells (Fig. 1a). In order to generate a functional, matrix-free bone marrow organoid from human iPSCs, we developed a simplified two-step protocol (Fig. 1b). iPSCs were aggregated into embryoid bodies (EBs) and induced toward a mesodermal lineage with the use of BMP4 and VEGF^16,17^, producing coral-like structures resembling early bone development. Subsequent floating culture supported further structural maturation and colonization by hematopoietic progenitors. Knockout serum replacement-based medium supported organoid formation across multiple iPSC lines. By day 3, organoids displayed irregular morphologies with arm-like extensions that persisted through day 14 (Fig. 1c and Supplementary Fig. 1a), mimicking elongation seen during long bone morphogenesis. Two independent iPSC lines produced similar structures.

**Fig. 1.**
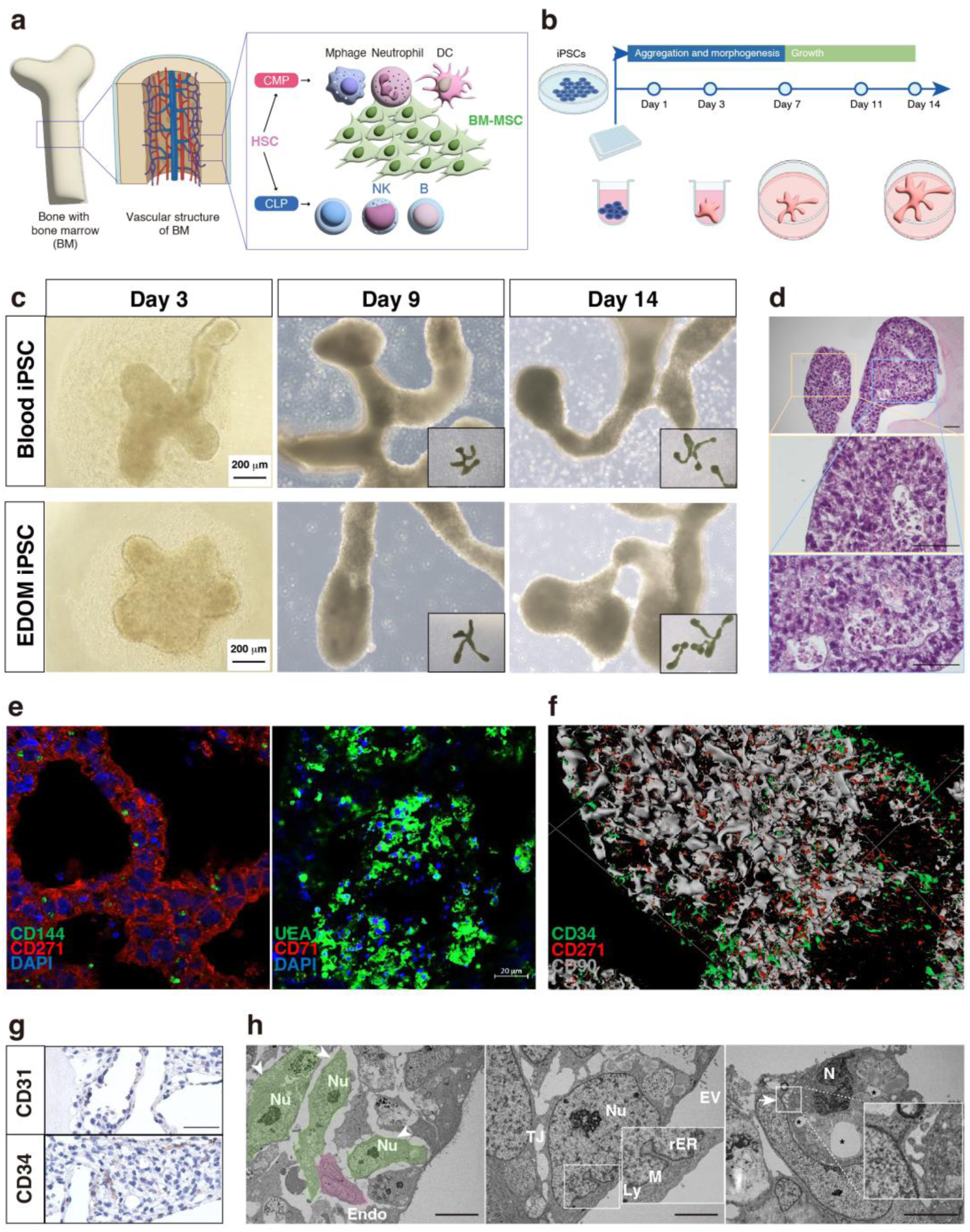
Generation and characterization of induced bone marrow organoids (iBMOs) from human iPSCs. **a.** Schematic of human bone marrow (BM) architecture highlighting vascular structures and key cell types, including myeloid, lymphoid, endothelial, and mesenchymal stromal cells. The BM is a semi-solid organ containing specialized vascular niches that support hematopoiesis. **b.** Workflow for Ibmo induction under matrix-free, self-organizing conditions. Aggregated embryoid bodies were induced toward mesoderm with BMP4 and VEGF, followed by floating culture to promote vascularization and hematopoietic stem/progenitor cell (HSPC) colonization. **c.** iBMOs exhibited a coral-like morphology during culture, consistently observed across two independent iPSC lines. **d.** Hematoxylin and eosin staining of day 14 iBMOs showed organized tissue architecture with vascular-like structures. Scale bars: 50 μm. **e** Immunofluorescence staining for VE-cadherin (CD144, endothelial marker), CD271 (NGFR1, stromal marker), UEA-I (endothelial marker), and CD71 (erythroid marker) revealed spatial organization of vascular and hematopoietic elements. **f** Three-dimensional reconstruction of whole-mount immunostained iBMOs cleared with CUBIC, showing mesenchymal stromal cells (CD271, CD90) and HSPCs (CD34). **g** Immunohistochemistry for CD31 (endothelial) and CD34 (HSPC) confirmed co-localization within iBMOs. Scale bars: 50 μm. **h.** Transmission electron microscopy revealed stromal cells with nucleoli (Nu), endothelial cells (left), stromal cells with tight junctions (TJ), mitochondria (M), lysosomes (Ly), and extracellular vesicles (EV) (center), and macrophage-like cells with phagocytic vacuoles (asterisks), pseudopodia (arrows), and characteristic nuclear morphology (N) (right). Scale bars: 10 μm (left), 1 μm (center), 5 μm (right). NK, natural killer cell; DC, dendritic cell; HSC, hematopoietic stem cell; MSC, mesenchymal stromal cell.

Hematoxylin and eosin staining of day 14 iBMOs revealed vascular-like networks interspersed with cell colonies (Fig. 1d and Supplementary Fig. 1b). Immunostaining confirmed VE-cadherin (CD144) and CD271 (NGFR1) expression in stromal and endothelial compartments (Fig. 1e, left), as well as limited CD34 expression in hematopoietic progenitors. Lectin UEA-I staining further indicated sinusoidal endothelial organization (Fig. 1e, right)^16^. Whole-mount CUBIC clearing^18^ and staining for CD34, CD90, and CD271 revealed CD34⁺ cells encased by stromal networks resembling native niches (Fig. 1f). Immunohistochemistry confirmed the presence of CD31⁺ endothelial and CD34⁺ progenitor cells (Fig. 1g).

Transmission electron microscopy (TEM) revealed the ultrastructural features of mesenchymal stromal cells—irregular nuclei with nucleoli, tight junctions, mitochondria, lysosomes, and extracellular vesicles^19^—alongside macrophage-like cells with pseudopodia and phagocytic vacuoles (Fig. 1h and Supplementary Fig. 1c). These findings demonstrated that iBMOs self-organized into three-dimensional (3D), vascularized structures with stromal and hematopoietic elements, faithfully recapitulating the architecture of native bone marrow and providing a reproducible human platform for studying hematopoiesis.

### Transcriptional profile of iBMOs

We compared the transcriptional profile of day 14 iBMOs with those of adult human bone marrow and progenitor iPSCs using bulk RNA sequencing. Hierarchical clustering showed that iBMOs closely resembled adult bone marrow and were clearly distinct from iPSCs (Supplementary Fig. 2). Extracellular matrix-related genes (COL3A1, LUM) and stromal-associated genes (IGFBP7, SLPI) were enriched in both the iBMOs and bone marrow, whereas pluripotency genes were downregulated, thus confirming lineage commitment.

Single-cell RNA sequencing (scRNA-seq) of 63,935 cells from day 14 iBMOs resulted in 12 transcriptionally distinct clusters (Fig. 2a). NCAM1 (CD56) expression was similar to that of PDGFRA, a stromal marker (Fig. 2b). Stromal populations expressed canonical markers (PDGFRA, COL1A1, COL3A1, IGFBP7), while endothelial clusters expressed VEGFC and APLNR, consistent with histological findings (Fig. 2c).

**Fig. 2.**
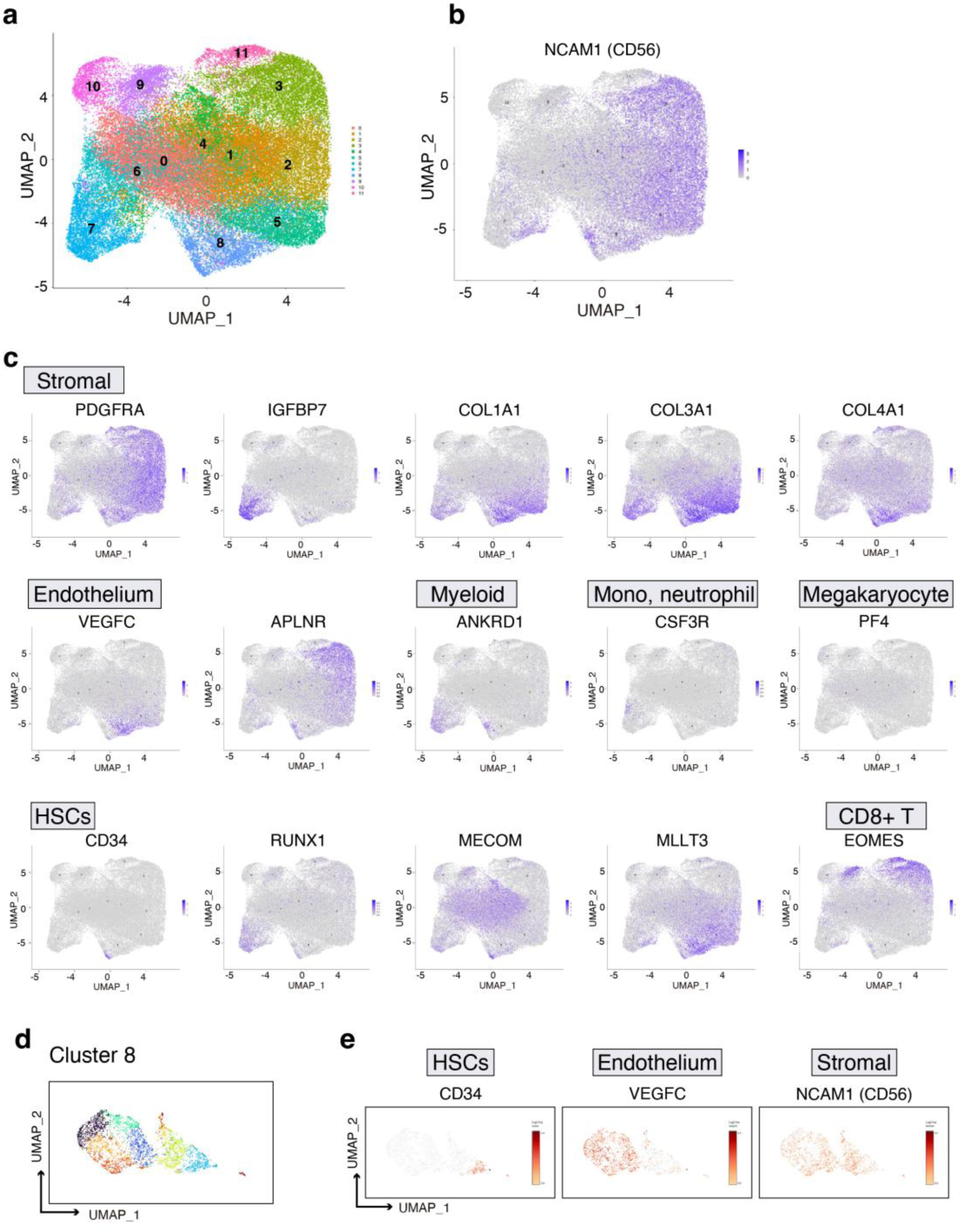
Single-cell RNA sequencing (scRNA-seq) analysis of iBMOs from human iPSCs. **a.** Uniform Manifold Approximation and Projection (UMAP) plots of integrated scRNA-seq data from four independent iBMO samples (63,935 cells) revealed 12 transcriptionally distinct clusters. **b.** NCAM1 (CD56) expression overlapped with PDGFRA, indicating stromal identity. **c.** Lineage markers detected included stromal markers (PDGFRA, IGFBP7, COL1A1, COL3A1, COL4A1), endothelial markers (VEGFC, APLNR), and a myeloid marker (ANKRD1). Hematopoietic clusters comprised monocytes (CSF3R), megakaryocytes (PF4), and CD8⁺ T lineage progenitors (EOMES), as well as hematopoietic stem/progenitor cell (HSPC) markers (CD34, RUNX1, MECOM, MLLT3). **d.** Cluster 8 was enriched for features of the hematopoietic niche. **e.** CD34 expression was localized to a discrete subregion, while VEGFC (endothelial) and NCAM1 (stromal) were expressed in adjacent regions within the same cluster, indicating a niche-like architecture with integrated vascular, stromal, and hematopoietic progenitor elements. HSC, hematopoietic stem cell.

Hematopoietic clusters included monocytes (CSF3R), megakaryocytes (PF4), and myeloid cells (ANKRD1). A hematopoietic stem/progenitor cell (HSPC) population co-expressed CD34, RUNX1, MECOM, and MLLT3, a factor essential for primitive HSC maintenance^20^, indicating bona fide HSC-like cells. A distinct EOMES⁺ cluster suggested the presence of CD8⁺ T lineage progenitors^21^.

Cluster-level mapping identified a unique subset (Cluster 8) co-expressing HSPC (CD34), endothelial (VEGFC), and stromal (NCAM1) markers (Fig. 2d, e). This hybrid signature suggests a specialized in vitro niche where stem, stromal, and vascular elements converge, resembling in vivo bone marrow microenvironments.

### Derivation and characterization of iBOSS

scRNA-seq of iBMOs revealed a predominant stromal population marked by NCAM1 (CD56), a feature shared by bone marrow stromal cells that contribute to hematopoietic niche formation^22^. Flow cytometry confirmed a substantial CD56⁺CD45⁻ fraction (Fig. 3a) co-expressing CD90 (THY-1), CD271 (NGFR1), and CD140a (PDGFRA) (Fig. 3b).

**Fig. 3.**
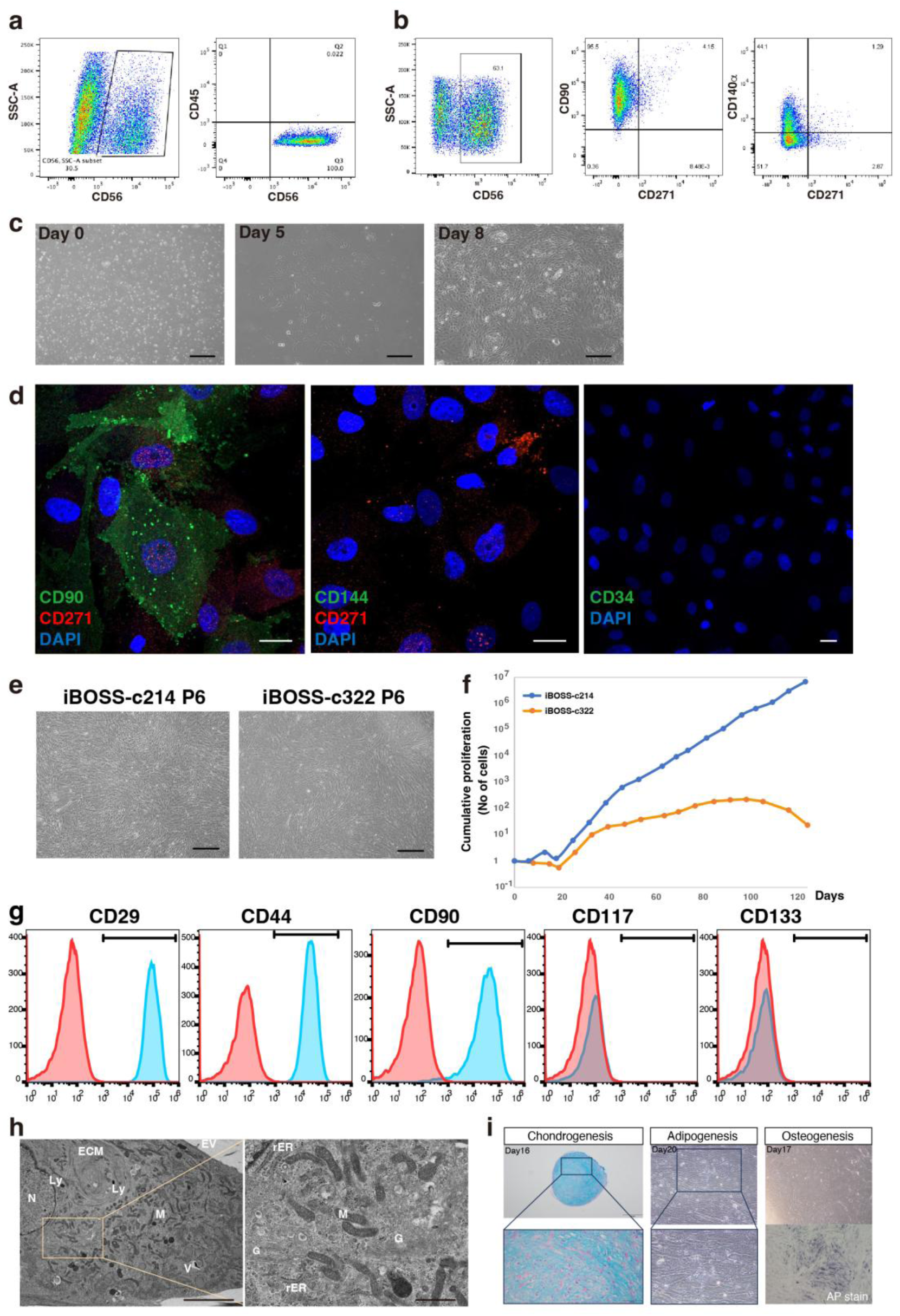
Establishment and characterization of iBMO-derived stromal stem cells (iBOSS). **a.** Flow cytometry revealed a substantial CD56⁺ population in iBMOs lacking the pan-hematopoietic marker CD45. **b.** CD56⁺ cells co-expressed bone marrow stromal markers CD90 (THY-1), CD271 (NGFR1), and CD140a (PDGFRA). **c.** Phase-contrast images of dissociated iBMO cells cultured without extracellular matrix coating on days 0 (left), 5 (middle), and 8 (right). Scale bars: 500 μm (left), 200 μm (middle and right). **d.** Immunofluorescence staining of adherent cells confirmed stromal identity: CD90 (green) and CD271 (red) (left); CD144 (green) and CD271 (red) (middle); CD34 (green) (right). Scale bar: 20 μm. **e.** Phase-contrast images of two iBOSS lines (c214 and c322) at passage 6, showing adherent spindle-shaped cells. Scale bar: 500 μm. **f.** Growth curves of iBOSS lines demonstrated clone-dependent proliferation rates. **g.** Flow cytometry confirmed expression of mesenchymal markers CD29, CD44, and CD90, with absence of hematopoietic markers. **h.** Transmission electron microscopy revealed mesenchymal ultrastructures, including collagen fibers, mitochondria (M), lysosomes (Ly), vacuoles (V), extracellular vesicles (EV), Golgi complexes (G), and rough endoplasmic reticulum (rER). Scale bars: 5 μm (left), 1 μm (right). **i.** iBOSS exhibited trilineage differentiation capacity, forming chondrocytes (Alcian Blue, left), adipocytes (lipid accumulation, center), and osteoblasts (AP stain; alkaline phosphatase activity, right).

When plated on uncoated tissue culture plastic, these cells adhered, proliferated, and yielded spindle-shaped cells (Fig. 3c, d). Non-adherent hematopoietic (CD34⁺) and endothelial (CD144⁺) cells were lost during passaging. The resulting cell lines, termed iBMO-derived stromal stem cells (iBOSS), proliferated stably with clone-dependent growth differences (Fig. 3e, f). Flow cytometry confirmed MSC markers (CD29, CD44, CD90) and absence of hematopoietic progenitor markers (CD117, CD133) (Fig. 3g). TEM revealed abundant rough ER, Golgi complexes, mitochondria, lysosomes, and extracellular matrix fibers (Fig. 3h.

Functionally, iBOSS showed multilineage mesenchymal differentiation: chondrogenic pellets (Alcian Blue⁺), adipocytes with lipid droplets, and osteoblasts with alkaline phosphatase activity (Fig. 3i). Collectively, iBOSS meet MSCs criteria, can be robustly derived from iPSCs, and offer a standardized human feeder platform. They provide an alternative to murine OP9 cells, which lack macrophage colony–stimulating factor^23–25^, and establish a human-compatible feeder system for scalable hematopoietic differentiation and screening applications. The ability to derive iBOSS lines reproducibly from multiple iPSC clones highlights their applicability in large-scale, xeno-free hematopoietic cell production systems.

### iBOSS drive T cell and DC differentiation

Notch ligands DLL4 and VCAM1 are critical for thymic T cell differentiation^26,27^, while Notch also influences DC and macrophage fates. scRNA-seq showed iBOSS expressed NOTCH1–3 and elevated DLL3 and VCAM1 compared with heterogeneous iBMO populations, which displayed high DLL1, restricted DLL3, and low DLL4 and VCAM1 (Fig. 4a and Supplementary Fig. 3). These findings suggest iBOSS provide an integrated microenvironment for Notch-dependent hematopoietic development.

**Fig. 4.**
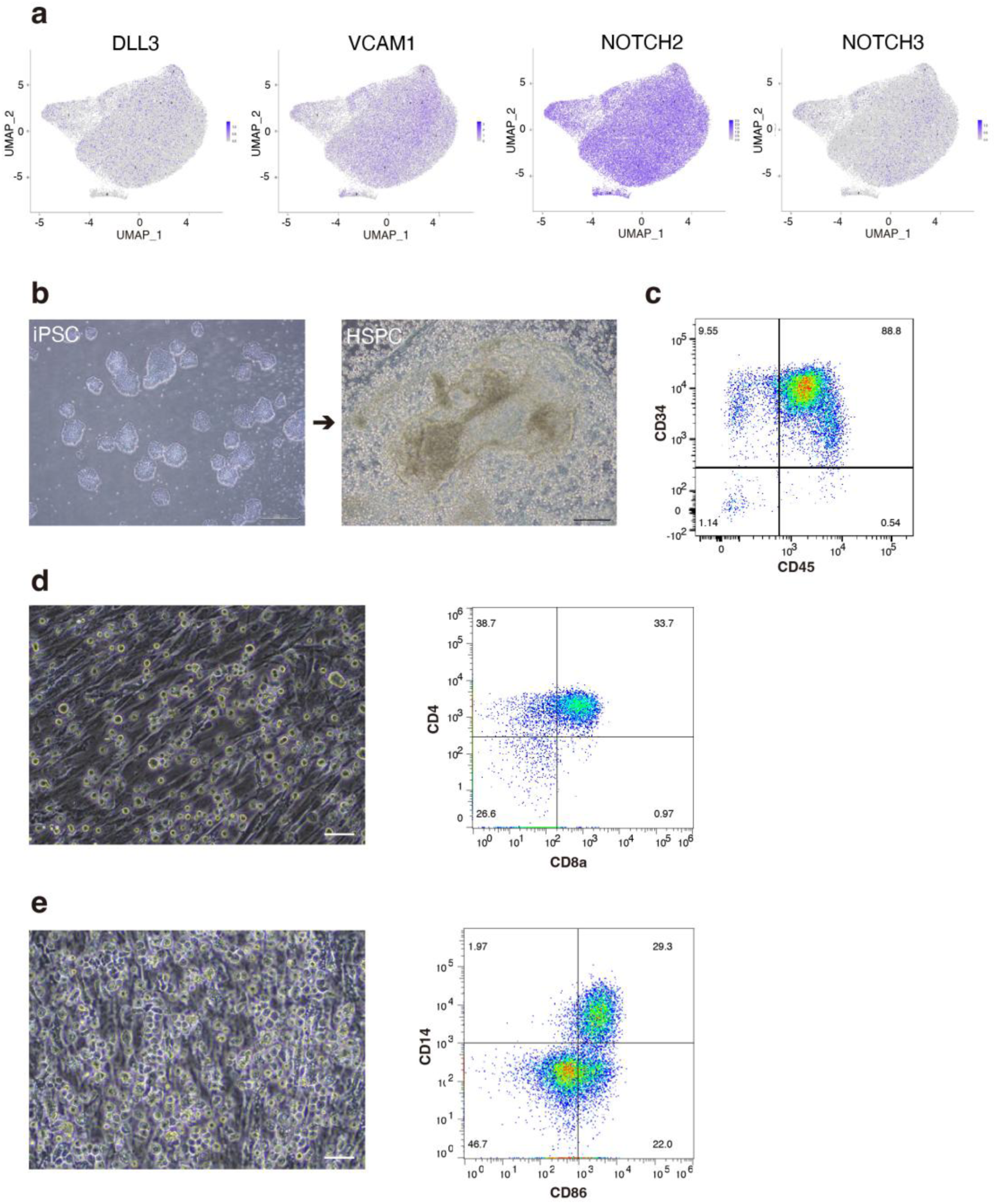
In vitro differentiation of T cells and dendritic cells (DCs) via co-culture of iPSC-derived hematopoietic progenitors with iBOSS. **a.** scRNA-seq analysis of iBOSS mesenchymal stromal cells. Integrated data from four iBOSS samples (54,084 cells) showed UMAP expression of DLL3, VCAM1, NOTCH2, and NOTCH3. **b.** Generation of hematopoietic stem/progenitor cells (HSPCs) from human iPSCs. Phase-contrast images show iPSCs before induction (left) and released HSPCs after 14 days of culture (right). Scale bars: 500 μm (left), 200 μm (right). **c** Flow cytometry confirmed released HSPCs were CD34⁺CD45⁺ double-positive. **d.** Flow cytometry revealed that CD34⁺ HSPCs differentiated into T cells with CD4 and CD8a expression in the presence of IL-7 on iBOSS. After 13 days, CD45⁺ cells comprised 38.7% CD4 single-positive and 33.7% CD4/CD8a double-positive populations. Scale bar: 50 μm. **e.** Flow cytometry showed that CD34⁺ HSPCs differentiated into DCs with CD86 expression in the presence of IL-4 and GM-CSF on iBOSS. At day 13, CD45⁺ cells included 2.7% CD14 single-positive and 40.9% CD86⁺ populations. Scale bar: 50 μm.

Human iPSC-derived HSPCs were generated via EB differentiation with TPO, SCF, and FLT3^17^, yielding a CD34⁺ cells (Fig. 4b, c). In the presence of IL-7, essential for thymic progenitor^28,29^, co-culture of HSPCs on iBOSS induced T cell differentiation. After 13 days, flow cytometry identified CD4 single-positive and CD4/CD8 double-positive cells (Fig. 4d). Giemsa staining showed lymphocyte morphology, and immunostaining confirmed CD4⁺ cells (Supplementary Fig. 4a), consistent with thymic-like induction^30,31^.

For DC induction^32,33^, HSPCs co-cultured on iBOSS with GM-CSF and IL-4^34,35^ produced adherent cells with stellate morphology (Fig. 4e, left). Flow cytometry identified CD86^⁺^CD45^⁺^ subsets, including CD14^⁺^/CD86^⁺^ and CD14^⁻^/CD86^⁺^ cells (Fig. 4e, right). DC identity was further confirmed by HLA-DR and CD86 staining (Supplementary Fig. 4b). Thus, iBOSS support the differentiation of iPSC-derived progenitors into both T cells and DCs, recapitulating thymic and marrow stromal functions in a single system.

### In vivo hematopoietic and ossifying potential of iBMOs

Day 14 iBMOs were transplanted into immunodeficient NOG mice^36^ under the renal capsule (RC) or subcutaneously (Fig. 5a and Supplementary Fig. 5a), where both ectopic sites lacked hematopoietic or osteogenic niches^37^. After six weeks, grafts at both sites expanded substantially, with RC implants showing more pronounced growth (Fig. 5b and Supplementary Fig. 5b). Histology revealed organized hematopoietic regions containing megakaryocyte-like cells and erythroblasts at multiple maturation stages, including perinuclear halo-bearing erythroid cells (Fig. 5c, left and center; and Supplementary Fig. 5c). Human origin was confirmed by LAMIN A/C and glycophorin A (CD235a) staining (Supplementary Fig. 5d). RC-engrafted iBMOs also underwent endochondral ossification, producing cartilage, cortical bone, and a periosteum-like layer (Fig. 5d).

**Fig. 5.**
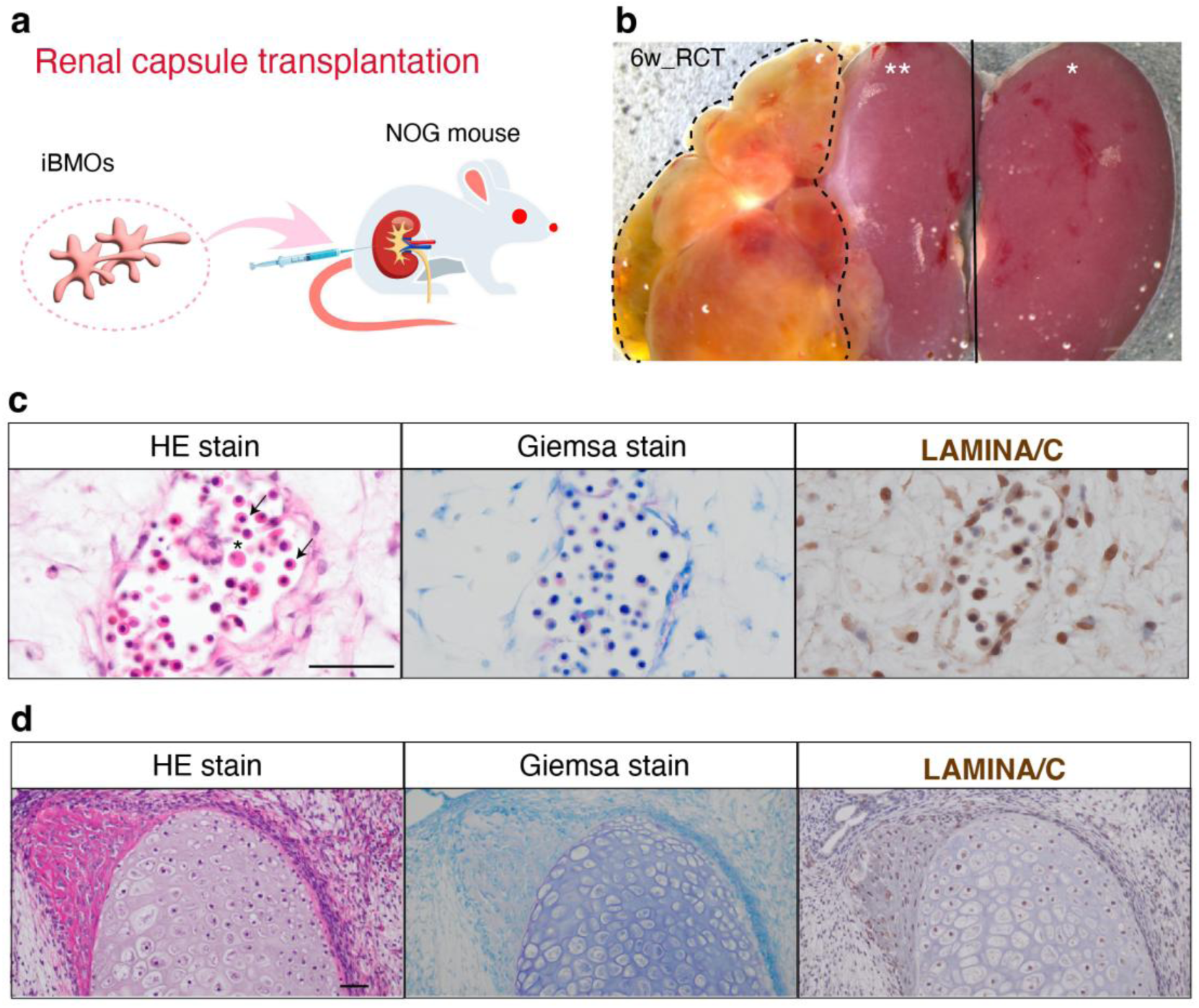
In vivo hematopoietic and ossifying potential of human iBMOs. **a.** Schematic of the transplantation strategy. Day 14 iBMOs were engrafted into immunodeficient NOG mice under the renal capsule. **b.** Representative images of explanted grafts six weeks post-transplantation. Renal capsule (RC) grafts showed marked enlargement and advanced maturation compared with subcutaneous grafts. **c.** Histology of RC-engrafted iBMOs. Hematoxylin and eosin (left) and Giemsa (center) staining revealed active erythropoiesis, including megakaryocyte-like cells (asterisks) and immature erythrocytes. Arrows indicate haloed perinuclei characteristic of erythroid maturation. Immunohistochemistry for human-specific nuclear marker LAMIN A/C confirmed human origin (right). **d.** Engrafted iBMOs also exhibited bone-forming potential. Hematoxylin and eosin (left) and Giemsa (center) staining showed cartilage and cortical bone, and LAMIN A/C staining confirmed human origin of bone-forming cells (right). Scale bars: 50 μm (C, D). RC, renal capsule.

Even at ectopic sites lacking supportive cues, iBMOs established a marrow-like niche, sustained human erythropoiesis, and generated mineralized bone, demonstrating integrated hematopoietic and skeletal potential *in vivo*, and establishing a foundation for humanized models of hematopoiesis and bone regeneration. These findings demonstrate that iBMOs can serve as a transplantable, self-contained human marrow analog, bridging in vitro modeling and in vivo regenerative applications.

## Discussion

In this study, we generated iBMOs from human iPSCs that recapitulated key structural and cellular features of the native bone marrow niche. Remarkably, these organoids also supported thymus-like T lymphopoiesis. We established a novel stromal stem cell line, iBOSS, as the source of this thymic-like activity. Together, these systems provide an integrated and defined, xeno-free *in vitro* platform that models fundamental aspects of bone marrow and thymic biology, creating new opportunities to study hematopoiesis and advance regenerative medicine, including applications in disease modeling and drug development.

iBMOs self-organized into complex 3D structures containing hematopoietic progenitors, vascular networks, and stromal cells, closely mirroring native bone marrow at histological and transcriptomic levels. When engrafted into immunodeficient mice, iBMOs autonomously supported human erythropoiesis and bone formation at ectopic sites. They therefore represent bona fide organoids with self-contained hematopoietic niches, independent of host microenvironments. This advancement surpasses prior bone marrow-like models, which have not shown comparable in vivo potential^16^, and validates our simplified, matrix-free culture system for generating highly functional organoids.

A key and unexpected finding was the capacity of iBOSS cells to support the differentiation of HSPCs into CD4⁺ single-positive and CD4⁺/CD8⁺ double-positive T cells, suggesting that bone marrow-derived stromal systems can partially recapitulate thymic cues for T cell development^8^. This activity appears to be mediated, at least in part, by Notch pathway components such as DLL3 and VCAM1 in iBOSS cells. These findings raise fundamental questions about the latent developmental plasticity of bone marrow stromal populations. Because iBOSS lines are genetically stable and human-derived, they offer a valuable alternative to murine feeder systems such as OP9 cells^23,24^, which lack supportive capacity for T cells.

Beyond T cells, the iBOSS co-culture system also generated functional DCs, enabling simultaneous derivation of lymphoid and myeloid lineages from a single iPSC-derived platform. This “all-in-one” approach provides a powerful model for dissecting lympho–myeloid interactions and could help meet the growing demand for large, standardized supplies of immune cells for adoptive cell therapy. The system also provides a donor-independent source for applications such as mixed lymphocyte reactions to assess transplant compatibility^38^. Moreover, because both iBMOs and iBOSS can be derived reproducibly from multiple iPSC lines, this approach provides a scalable, donor-independent framework for manufacturing patient-specific immune cells under defined conditions.

There are limitations to this study. The T cell populations generated here were not fully characterized to mature cytotoxic or helper subsets, and their functional competence was not assessed. Future studies are required to determine their degree of maturation and antigen specificity.

In conclusion, the iBMO–iBOSS platform functionally integrates essential features of bone marrow and thymic niches, enabling the coordinated generation of multiple hematopoietic lineages from human iPSCs. This system provides a unique opportunity to probe principles of human hematopoietic development and may serve as a reproducible and scalable source of immune cells for translational immunotherapy. By unifying bone marrow and thymic functions in a single human-derived model, this work expands our understanding of human hematopoietic biology and opens new avenues for engineering synthetic, patient-specific immune systems.

## Methods

### Human induced pluripotent stem cell (iPSC) culture

At least three human iPSC lines were used and cultured in StemFlex complete medium (StemFlex Basal Medium [A33493-01, Gibco] supplemented with StemFlex Supplement [A33492-01, Gibco]). For passaging, dissociated human iPSCs were seeded into rhVTN-N (A14700, Gibco)-coated 6-well plates with 10 µM Rock inhibitor (Y27632) in StemFlex complete medium and incubated at 37 °C with 5% CO_2_. The next day, the medium was fully changed to remove the Rock inhibitor. Cells were cultured for up to one week with medium changes every 1–3 days before the next passage.

### Generation of induced bone marrow organoids (iBMOs)

Dissociated iPSCs were seeded into U-bottom 96-well plates (174929, Thermo Fisher) in bone marrow basal medium (BMBM), composed of StemPro34 complete medium (10639-011, Gibco), 20% KSR (10828-028, Gibco), penicillin/streptomycin, monothioglycerol (MTG, M1753-100ML, Sigma), GlutaMAX, L-ascorbic acid, and ITS-A (51300044, Gibco), supplemented with BMP4, bFGF, VEGF, and Y27632. After 1 day, the medium was changed to remove Y27632. On day 3, the medium was replaced with BMBM supplemented with BMP4 (Qk038, Qkine), bFGF, VEGF (Qk048, Qkine), SCF (255-SC-050, R&D Systems), and FLT3 ligand (308-FK-025, R&D Systems). On day 7, coral-like aggregates were transferred to larger dishes and maintained with the same cytokines. Organoids were collected after 14–17 days for further analysis or derivation of stromal cell lines.

### Establishment of iBMO-derived stromal stem cells (iBOSS)

Organoids were harvested on day 17 and dissociated enzymatically using collagenase I and dispase II at 37°C for 20 min, followed by TrypLE Select (12563-011, Gibco) treatment for 5–10 min. Dissociated cells were plated on uncoated dishes in modified B0 medium (KnockOut DMEM [10829-018, Gibco] and Ham’s F-12 [11765054, Gibco] in a 2:1 ratio, 10% human serum, penicillin/streptomycin, MTG, L-ascorbic acid, and ITS-A), and cultured at 37 °C with 5% CO_2_.

### Immunohistochemistry and Immunofluorescence

Organoids that had been cultured for 14 days were fixed in 4% paraformaldehyde (PFA) in PBS and processed for paraffin embedding. Sections (5 µm thickness) were cut using a microtome, deparaffinized, and rehydrated. For antigen retrieval, sections were subjected to heat-induced epitope retrieval in Histofine antigen retrieval buffer using a microwave. Sections were then blocked and incubated with primary antibodies overnight at 4°C. After sections were washed with PBS, they were incubated with corresponding secondary antibodies for 1–2 hours at room temperature. Slides were then counterstained with DAPI and mounted.

The following primary antibodies were used: anti-VE-cadherin (CD144; MA1-198, Thermo Fisher Scientific), anti-CD271 (NGFR; MA5-31968, Thermo Fisher Scientific), anti-CD34 (343502, BioLegend), anti-CD71 (14-0719-82, Thermo Fisher Scientific), biotinylated Ulex Europaeus Agglutinin I (UEA-I; B-1065-2, Vector Laboratories), and anti-CD90 (Thy1; 328102, BioLegend). The following secondary antibodies and reagents were used: Alexa Fluor 568 goat anti-mouse IgG1 (A11004, Invitrogen), Alexa Fluor 568 goat anti-rabbit IgG (A11036, Invitrogen), Alexa Fluor 647 goat anti-mouse IgG (A32728, Invitrogen), Alexa Fluor 488 goat anti-mouse IgG (A28175, Invitrogen), and Streptavidin-Alexa Fluor 488 (S11223, Invitrogen). For whole-mount analysis, day 14 iBMOs were fixed with 4% PFA in PBS. Tissue clearing was performed using CUBIC-L and CUBIC-R solutions (TCI) according to the manufacturer’s instructions. Cleared organoids were then immunostained with the following fluorescent conjugated primary antibodies: Alexa Fluor 647 anti-CD90/Thy1 (ab202334, Abcam), Alexa Fluor 488 anti-CD34 (ab187568, Abcam), and Alexa Fluor 594 anti-NGFR (NBP2-47966AF594, Novus Biologicals). Stained iBMOs were imaged using a Zeiss LSM 900 laser scanning confocal microscope.

### RNA sequencing analysis

Organoids were collected after 14 days of culture (n = 8) and immediately frozen in liquid nitrogen. Human iPSCs (two independent lines) and commercially obtained human bone marrow RNA were used as controls. Total RNA was extracted using the miRNeasy Micro Kit (217084, QIAGEN) following the manufacturer’s instructions. RNA sequencing libraries were prepared using the TruSeq Stranded mRNA Library Prep Kit (Illumina) and sequenced on NovaSeq 6000 and NovaSeq X platforms (paired-end, 100 bp). Sequencing reads were aligned to the human reference genome (hg38), and transcript abundance was calculated in transcripts per million (TPM) for each gene.

### Single-cell RNA sequencing (scRNA-seq) analysis

For scRNA-seq of iBMOs, day 14 cultured iBMOs (n = 4 samples, derived from two iPSC lines; 16 organoids per sample) were dissociated into single cells using enzymatic digestion with Dispase II and collagenase I, followed by two rounds of filtration through 40 μm cell strainers. For iBOSS samples (n = 4 from three iPSC lines), monolayer cells were dissociated using TrypLE Select. Cell viability (>85%) was confirmed using the ReadyCount Green/Red Viability Stain (A49905, Invitrogen) and measured with a Countess 3 FL automated cell counter (Invitrogen).

Single-cell libraries were prepared using the Chromium Next GEM Single Cell 3′ Kit v3.1 (Dual Index, 10x Genomics) according to the manufacturer’s instructions. Approximately 16,500 cells per sample were loaded into the Chromium Controller (10x Genomics) with reagents and barcoded gel beads. Single-cell Gel Beads-in-Emulsion (GEMs) were generated by microfluidic partitioning, cell lysis enablement, reverse transcription, and incorporation of unique molecular identifiers (UMIs) and cell barcodes. After emulsion breaking, barcoded cDNA was purified using Dynabeads MyOne SILANE and amplified by PCR. The resulting cDNA was used to construct sequencing libraries with the Chromium Library Construction Kit, which was followed by size selection and purification. Sequencing was performed on an Illumina NovaSeq X Plus platform. Read configuration was as follows: Read 1, 28 bp (cell barcode and UMI); i7 and i5 indices, 10 bp each; Read 2, 90 bp (transcript). Target sequencing depth ranged from 25,523 to 50,871 reads per cell. Raw FASTQ files were processed using Cell Ranger (v7.0, 10x Genomics) to generate gene–cell UMI count matrices. Cells with fewer than 500 detected genes, more than 20% mitochondrial transcript content, or suspected doublets were excluded. Downstream normalization and batch correction were performed using the Seurat R package. scRNA-seq data from four iBMO samples (total cells = 63,935) and four iBOSS samples (total cells = 54,084) were integrated separately. For data visualization, Uniform Manifold Approximation and Projection (UMAP) plots were generated using the R platform or Loupe Browser v8 (10x Genomics).

### Flow cytometry analysis

For flow cytometry analysis, all samples were analyzed with a FACS Aria II (BD), SH800, or MA900 flow cytometer (Sony). iBMOs were collected after 14 days of culture, treated with collagenase I and dispase II at 37°C for 20 min, and then treated with diluted TrypLESelect at 37°C for 5–10 min. The iBMOs were separated into single cells by pipetting. The cells were centrifuged and resuspended with 2%FBS-PBS, and then stained with fluorescent conjugated antibodies: FITC anti-human CD45 Antibody (304005), PE anti-human CD45 Antibody (368509), APC anti-human CD56 (NCAM1) Antibody (362503), PE/ Cyanine7 anti-human CD56 (NCAM1) Antibody (362509), APC/Cyanine7 anti-human CD271 (NGFR) Antibody (345125), PE anti-human CD140a (PDGFRα) Antibody (323505), and Brilliant Violet 421 anti-human CD90 (Thy1) Antibody (328121, Biolegend for all antibodies). The hematopoietic stem/progenitor cells (HSPCs) derived from human iPSCs were collected after 14 days of culture and stained with fluorescent conjugated antibodies FITC anti-human CD45 Antibody and APC anti-human CD34 Antibody (378605, Biolegend) to confirm HSPC properties. For the mesenchymal stromal cell characterization of established iBOSS, flow cytometry analysis was performed by surface marker staining with antibodies: FITC anti mouse/human CD44 antibody (11-0441, eBio), Brilliant Violet 421 anti-human CD90 (Thy1) Antibody (328121, Biolegend), PE anti-human CD29 Antibody, (303003, Biolegend), PE anti-human CD117 (c-kit) Antibody (313203, Biolegend), and PE anti-human CD133 Antibody (397903, Biolegend). The following isotype control antibodies were used: FITC Rat IgG2b Isotype control (11-4031, eBio), Brilliant Violet 421 Mouse IgG1, κ Isotype Ctrl Antibody (400157, Biolegend), PE Mouse IgG1, and κ Isotype Ctrl (FC) Antibody (400113, Biolegend). T and dendritic cell (DC) differentiation was analyzed using specific antibodies: FITC anti-human CD45 Antibody, Brilliant Violet 421 anti-human CD4 Antibody (344631, Biolegend), Brilliant Violet 570 anti-human CD8a Antibody (301037, Biolegend), Brilliant Violet 421anti-human CD14 Antibody (325627, Biolegend), and PE anti-human CD86 Antibody (374205, Biolegend). Flow data were analyzed using FlowJo software when necessary.

### In vitro differentiation of iBOSS

The developmental potential of iBOSS cells toward mesenchymal lineages was assessed under defined culture conditions. Adipogenic, chondrogenic, and osteogenic differentiation processes were induced using commercially available kits (hMSC Adipogenic Differentiation BulletKit, PT-3004; Chondrogenic, PT-3003; Osteogenic, PT-3002; Lonza) according to the manufacturer’s instructions. After culture for 20 days (adipogenesis), 16 days (chondrogenesis), or 17 days (osteogenesis), Alcian blue staining was used to detect glycosaminoglycan-rich extracellular matrix in chondrocytes, while alkaline phosphatase activity was evaluated as a marker of osteogenic differentiation.

### Differentiation of human iPSC-derived hematopoietic progenitors into T cells and DCs using iBOSS feeders

HSPCs were derived from human iPSCs through embryoid body formation based on a previously described protocol^17^ with modifications. HSPCs collected after ≥18 days of culture were transferred onto confluent monolayers of iBOSS cells for lineage-specific differentiation.

Co-cultures were maintained in B0-modified medium (KnockOut DMEM: Ham’s F-12 = 2:1, supplemented with 10% human serum, penicillin-streptomycin, monothioglycerol, L-ascorbic acid, and ITS-A) with BMP4, VEGF, SCF, and FLT3 for five days. From day 6 onward, IL-7 and FLT3 were added for T cell differentiation, while GM-CSF and IL-4 were used for dendritic cell induction. Cells were harvested at defined time points (days 6, 10, and/or 13) for downstream analyses.

### In vivo engraftment of iBMOs into immunodeficient mice

Day 14 cultured iBMOs were transplanted into immunodeficient NOG mice (NOD.Cg-Prkdc/ShiJic scidIl2rg tm1Sug3) to evaluate their in vivo developmental potential. All animal procedures were approved by the Animal Care and Use Committee of the National Center for Child Health and Development (A2003-002) and conducted in accordance with the 3Rs principle (refine, reduce, replace). All efforts were made to minimize animal suffering and reduce the number of animals used. The iBMOs were engrafted either subcutaneously or under the renal capsule using a 21-gauge needle. At six weeks post-transplantation, grafts were harvested, fixed in 20% neutral buffered formalin, and processed for histological examination. Hematoxylin and eosin and Giemsa staining were employed to assess hematopoietic and skeletal differentiation. The human origin of the cells was confirmed by immunohistochemistry for LAMIN A/C. Erythropoiesis within grafts was evaluated by immunofluorescence staining for human CD235a (Glycophorin A, ab129024, Abcam).

### Data and materials availability

All data are available in the main text or the supplementary materials. Bulk and single-cell RNA sequencing data have been deposited in GEO under accession number [to be provided upon availability]. Materials used in this study, including iPSC lines and iBOSS cell lines, are available from the corresponding author upon reasonable request and may be subject to a materials transfer agreement (MTA).

## Acknowledgements

We thank M. Ichinose for providing expert technical assistance.

## Funding

This work was supported by the Japan Agency for Medical Research and Development (AMED) grant awarded to H.A. (grant nos. 24be0704001h0005, 24bm1323001h0302), the National Center for Child Health and Development grant awarded to H.A. (grant no. 2022A-2) and to J.L. (grants no. 2024C-26 and 2025C-8).

## Author contributions

Conceptualization: J.L. and H.A.

Methodology: J.L.

Investigation: J.L., T.K., L.T., T.U.

Visualization: J.L.

Funding acquisition: J.L. and H.A.

Project administration: J.L., H.A.,A.U.

Writing – original draft: J.L. and H.A.

Writing – review & editing: J.L., H.A., A.U., S.Y.

## Competing interest declaration

NCCHD filed patent applications (JP2024-112515, PCT/JP2025/22044) on the content of the manuscript. J.L., H.A., and A.U. are the inventors of the patent.

## Notes

### Competing Interest Statement

The authors have declared no competing interest.

## References

1 Calvanese, V. & Mikkola, H. K. A. The genesis of human hematopoietic stem cells. Blood 142, 519–532 (2023). 10.1182/blood.2022017934

2 Laurenti, E. & Gottgens, B. From haematopoietic stem cells to complex differentiation landscapes. Nature 553, 418–426 (2018). 10.1038/nature25022

3 Mendez-Ferrer, S. et al. Mesenchymal and haematopoietic stem cells form a unique bone marrow niche. Nature 466, 829–834 (2010). 10.1038/nature09262

4 Sovani, V. Normal bone marrow, its structure and function. Diagnostic Histopathology 27, 349–356 (2021). 10.1016/j.mpdhp.2021.06.001

5 Herzog, E. L., Chai, L. & Krause, D. S. Plasticity of marrow-derived stem cells. Blood 102, 3483–3493 (2003). 10.1182/blood-2003-05-1664

6 Bianco, P., Robey, P. G. & Simmons, P. J. Mesenchymal stem cells: revisiting history, concepts, and assays. Cell Stem Cell 2, 313–319 (2008). 10.1016/j.stem.2008.03.002

7 Gao, Q. et al. Bone Marrow Mesenchymal Stromal Cells: Identification, Classification, and Differentiation. Front Cell Dev Biol 9, 787118 (2021). 10.3389/fcell.2021.787118

8 Schmitt, T. M. & Zuniga-Pflucker, J. C. Induction of T Cell Development from Hematopoietic Progenitor Cells by Delta-like-1 In Vitro. Immunity 17, 749–756 (2002).

9 Jones, R. J. Is post-transplant cyclophosphamide a true game-changer in allogeneic transplantation: The struggle to unlearn. Best Pract Res Clin Haematol 32, 101112 (2019). 10.1016/j.beha.2019.101112

10 Ciceri, F. et al. A survey of fully haploidentical hematopoietic stem cell transplantation in adults with high-risk acute leukemia: a risk factor analysis of outcomes for patients in remission at transplantation. Blood 112, 3574–3581 (2008). 10.1182/blood-2008-02-140095

11 Petersdorf, E. W. et al. Role of HLA-B exon 1 in graft-versus-host disease after unrelated haemopoietic cell transplantation: a retrospective cohort study. Lancet Haematol 7, e50–e60 (2020). 10.1016/S2352-3026(19)30208-X

12 Fabricius, W. A. & Ramanathan, M. Review on Haploidentical Hematopoietic Cell Transplantation in Patients with Hematologic Malignancies. Adv Hematol 2016, 5726132 (2016). 10.1155/2016/5726132

13 Bertaina, A. & Andreani, M. Major Histocompatibility Complex and Hematopoietic Stem Cell Transplantation: Beyond the Classical HLA Polymorphism. Int J Mol Sci 19, 621 (2018). 10.3390/ijms19020621

14 Huch, M. & Koo, B. K. Modeling mouse and human development using organoid cultures. Development 142, 3113–3125 (2015). 10.1242/dev.118570

15 Tsuruta, S., Uchida, H. & Akutsu, H. Intestinal Organoids Generated from Human Pluripotent Stem Cells. JMA J 3, 9–19 (2020). 10.31662/jmaj.2019-0027

16 Khan, A. O. et al. Human Bone Marrow Organoids for Disease Modeling, Discovery, and Validation of Therapeutic Targets in Hematologic Malignancies. Cancer Discov 13, 364–385 (2023). 10.1158/2159-8290.CD-22-0199

17 Iriguchi, S. et al. A clinically applicable and scalable method to regenerate T-cells from iPSCs for off-the-shelf T-cell immunotherapy. Nat Commun 12, 430 (2021). 10.1038/s41467-020-20658-3

18 Gomez-Gaviro, M. V. et al. Optimized CUBIC protocol for three-dimensional imaging of chicken embryos at single-cell resolution. Development 144, 2092–2097 (2017). 10.1242/dev.145805

19 Bauman, E., Feijao, T., Carvalho, D. T. O., Granja, P. L. & Barrias, C. C. Xeno-free pre-vascularized spheroids for therapeutic applications. Sci Rep 8, 230 (2018). 10.1038/s41598-017-18431-6

20 Calvanese, V. et al. MLLT3 governs human haematopoietic stem-cell self-renewal and engraftment. Nature 576, 281–286 (2019). 10.1038/s41586-019-1790-2

21 Kaech, S. M. & Cui, W. Transcriptional control of effector and memory CD8+ T cell differentiation. Nat Rev Immunol 12, 749–761 (2012). 10.1038/nri3307

22 Van Acker, H. H., Capsomidis, A., Smits, E. L. & Van Tendeloo, V. F. CD56 in the Immune System: More Than a Marker for Cytotoxicity? Front Immunol 8, 892 (2017). 10.3389/fimmu.2017.00892

23 Kodama, H., Nose, M., Niida, S., Nishikawa, S. & Nishikawa, S. Involvement of the c-kit receptor in the adhesion of hematopoietic stem cell to stromal cells. Exp Hematol 22, 979–984 (1994).

24 Schmitt, T. M. & Zuniga-Pflucker, J. C. T-cell development, doing it in a dish. Immunol Rev 209, 95–102 (2006). 10.1111/j.0105-2896.2006.00353.x

25 Nakano, T., Kodama, H. & Honjo, T. Generation of lymphohematopoietic cells from embryonic stem cells. Science 265, 1098–1101 (1994).

26 Zhou, B. et al. Notch signaling pathway: architecture, disease, and therapeutics. Signal Transduct Target Ther 7, 95 (2022). 10.1038/s41392-022-00934-y

27 Shukla, S. et al. Progenitor T-cell differentiation from hematopoietic stem cells using Delta-like-4 and VCAM-1. Nat Methods 14, 8 (2017). 10.1038/nmeth.4258532

28 Chen, D., Tang, T. X., Deng, H., Yang, X. P. & Tang, Z. H. Interleukin-7 Biology and Its Effects on Immune Cells: Mediator of Generation, Differentiation, Survival, and Homeostasis. Front Immunol 12, 747324 (2021). 10.3389/fimmu.2021.747324

29 Puel, A., Ziegler, S. F., Buckley, R. H. & Leonard, W. J. Defective IL7R expression in T-B+NK+ severe combined immunodeficiency. Nat Genet 20, 394–397 (1998).

30 Anderson, G., Moore, N. C., Owen, J. J. T. & Jenkinson, E. J. Cellular interactions in thymocyte development. Annu Rev Immunol 14, 27 (1996).

31 Hale, J. S. & Fink, P. J. Back to the thymus: peripheral T cells come home. Immunol Cell Biol 87, 58–64 (2009). 10.1038/icb.2008.87

32 Guilliams, M. et al. Dendritic cells, monocytes and macrophages: a unified nomenclature based on ontogeny. Nat Rev Immunol 14, 571–578 (2014). 10.1038/nri3712

33 Hart, D. N. J. Dendritic Cells: Unique Leukocyte Populations Which Control the Primary Immune Response. Blood 90, 3245–3287 (1997). 10.1182/blood.V90.9.3245

34 N’Diaye, M. et al. Rat bone marrow-derived dendritic cells generated with GM-CSF/IL-4 or FLT3L exhibit distinct phenotypical and functional characteristics. J Leukoc Biol 99, 437–446 (2016). 10.1189/jlb.1AB0914-433RR

35 Polancec, D. S. et al. Azithromycin drives in vitro GM-CSF/IL-4-induced differentiation of human blood monocytes toward dendritic-like cells with regulatory properties. J Leukoc Biol 91, 229–243 (2012). 10.1189/jlb.1210655

36 Shultz, L. D. et al. Human cancer growth and therapy in immunodeficient mouse models. Cold Spring Harb Protoc 2014, 694–708 (2014). 10.1101/pdb.top073585

37 Cunha, G. R. & Baskin, L. Use of sub-renal capsule transplantation in developmental biology. Differentiation 91, 4–9 (2016). 10.1016/j.diff.2015.10.007

38 Bromelow, K. V. et al. Whole blood assay for assessment of the mixed lymphocyte reaction. J Immunol Methods 247, 1–8 (2001).

