## Supplementary Figures for "Integration of Hematopoietic and Thymus-like Niches in a Human iPSC-derived Bone Marrow Organoid"

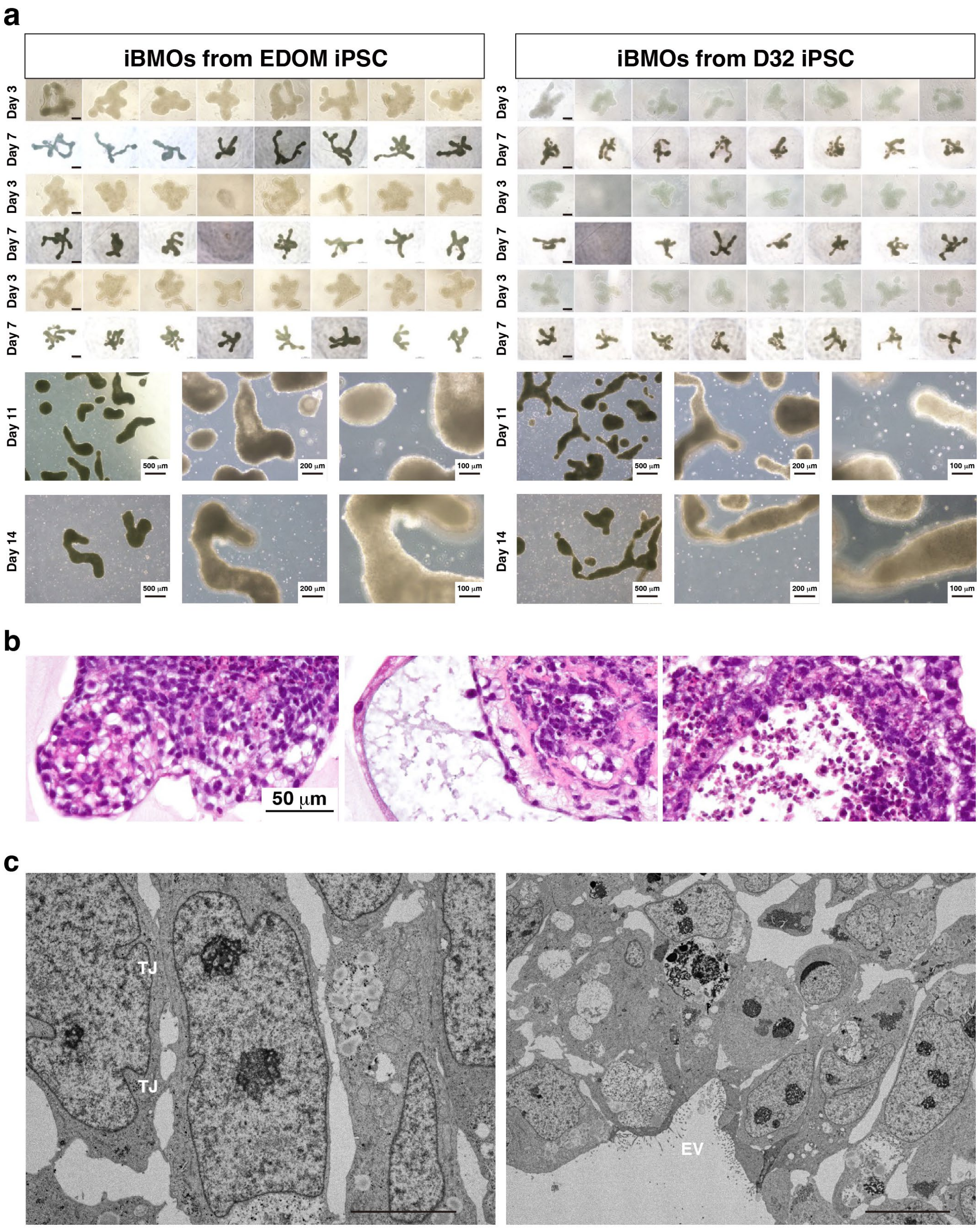


**Supplementary Figure 1. Generation and characterization of induced bone marrow organoids (iBMOs) from human iPSCs. a.** iBMOs from two independent iPSC lines (EDOM and D32) exhibited a characteristic coral-like morphology during in vitro culture. **b.** Hematoxylin and eosin (HE) staining of day 14 iBMOs revealed organized tissue architecture with vascular-like structures. **c.** Transmission electron microscopy showed stromal cells with tight junctions (TJ, left) and extracellular vesicles (EV, right). Scale bars: 5 μm (left), 10 μm (right).

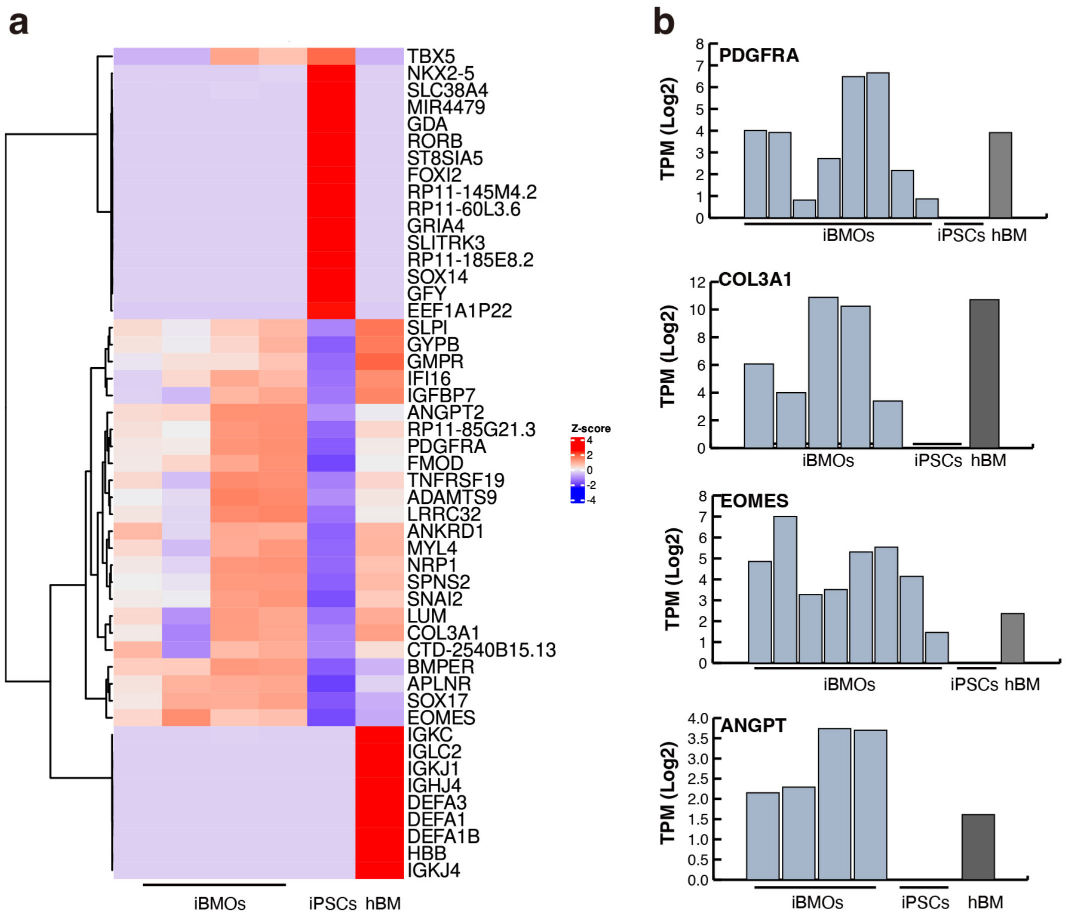


**Supplementary Figure 2. Transcriptional profile of day 14 iBMOs by RNA sequencing. a.** Heatmap and hierarchical clustering of day 14 iBMOs, adult human bone marrow (hBM), and progenitor iPSCs from bulk RNA sequencing. **b.** Gene expression levels of PDGFRA, COL3A1, EOMES, and ANGPT.

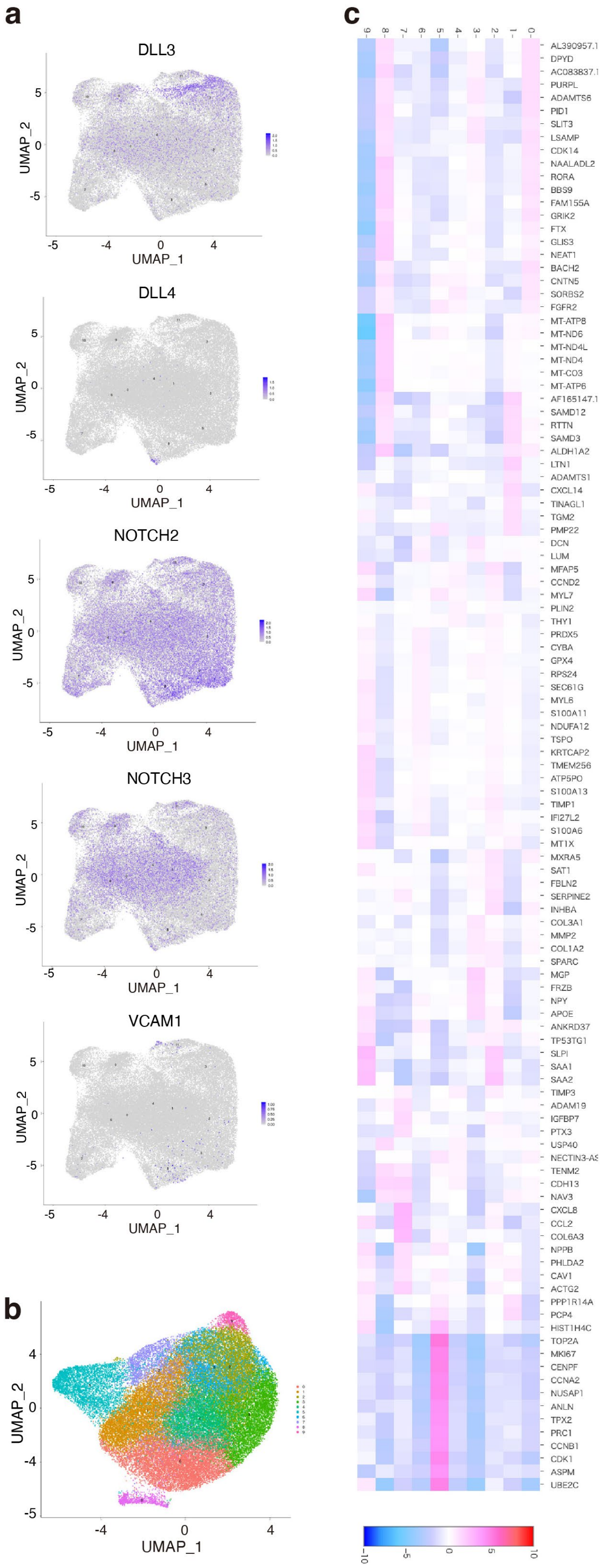

a
DLL3
DLL4
NOTCH2
NOTCH3
VCAM1
UMAP_1
UMAP_2
b
c
9 8 7 6 5 4 3 2 1 0
AL390957.1
DPYD
PURPL
ADAMTS6
PID1
SLIT3
LSAMP
CDK14
NAALADL2
RORA
BBS9
FAM155A
GRIK2
FTX
GLIS3
NEAT1
BACH2
CNTN5
SORBS2
FGFR2
MT-ATP8
MT-ND6
MT-ND4L
MT-ND4
MT-CO3
MT-ATP6
AF165147.1
SAMD12
RTTN
SAMD3
ALDH1A2
LTN1
ADAMTS1
CXCL14
TINAGL1
TGM2
PMP22
DCN
LUM
MFAP5
CCND2
MYL7
PLIN2
THY1
PRDX5
CYBA
GPX4
RPS24
SEC61G
MYL6
S100A11
NDUFA12
TSPO
KRTCAP2
TMEM256
ATP5PO
S100A13
TIMP1
IFI27L2
S100A6
MT1X
MXRA5
SAT1
FBLN2
SERPINE2
INHBA
COL3A1
MMP2
COL1A2
SPARC
MGP
FRZB
NPY
APOE
ANKRD37
TP53TG1
SLPI
SAA1
SAA2
TIMP3
ADAM19
IGFBP7
PTX3
USP40
TENM2
CDH13
NAV3
CXCL8
CCL2
COL6A3
NPPB
PHLDA2
CAV1
ACTG2
PPP1R14A
PCP4
HIST1H4C
TOP2A
MKI67
CENPF
CCNA2
NUSAP1
ANLN
TPX2
PRC1
CCNB1
CDK1
ASPM
UBE2C
-10
-5
0
5
10

**Supplementary Figure 3. Single-cell RNA sequencing (scRNA-seq) of iBMOs and iBMO and iBOSS. a.** UMAP plots of DLL3, DLL4, NOTCH2, NOTCH3, and VCAM1 in integrated data from four iBMO samples. **b.** UMAP plots of integrated data from four iBOSS samples (54,084 cells). **c.** Heatmap of differentially expressed genes among clusters identified in iBOSS.

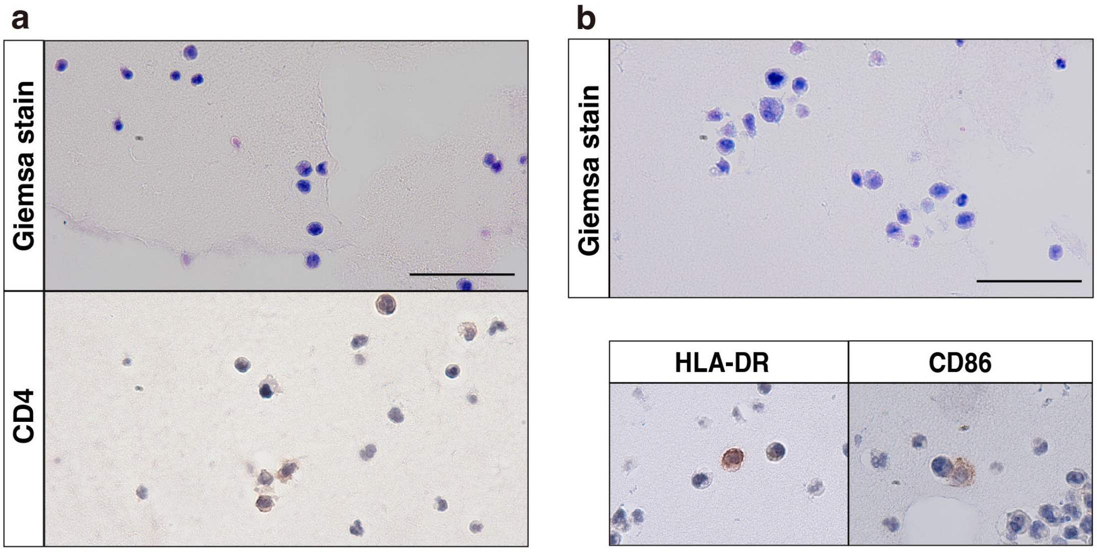


**Supplementary Figure 4. In vitro differentiation of T cells and dendritic cells (DCs) via co-culture of iPSC-derived HSPCs with iBOSS. a.** Giemsa staining revealed lymphocyte morphology with a dark purple nucleus and thin pink cytoplasm (top). Immunohistochemistry confirmed $CD4^+$ cells (bottom). **b.** Giemsa staining showed DC-like morphology (top), and immunohistochemistry demonstrated HLA-$DR^+$ and $CD86^+$ cells (bottom). Scale bars: 50 μm.

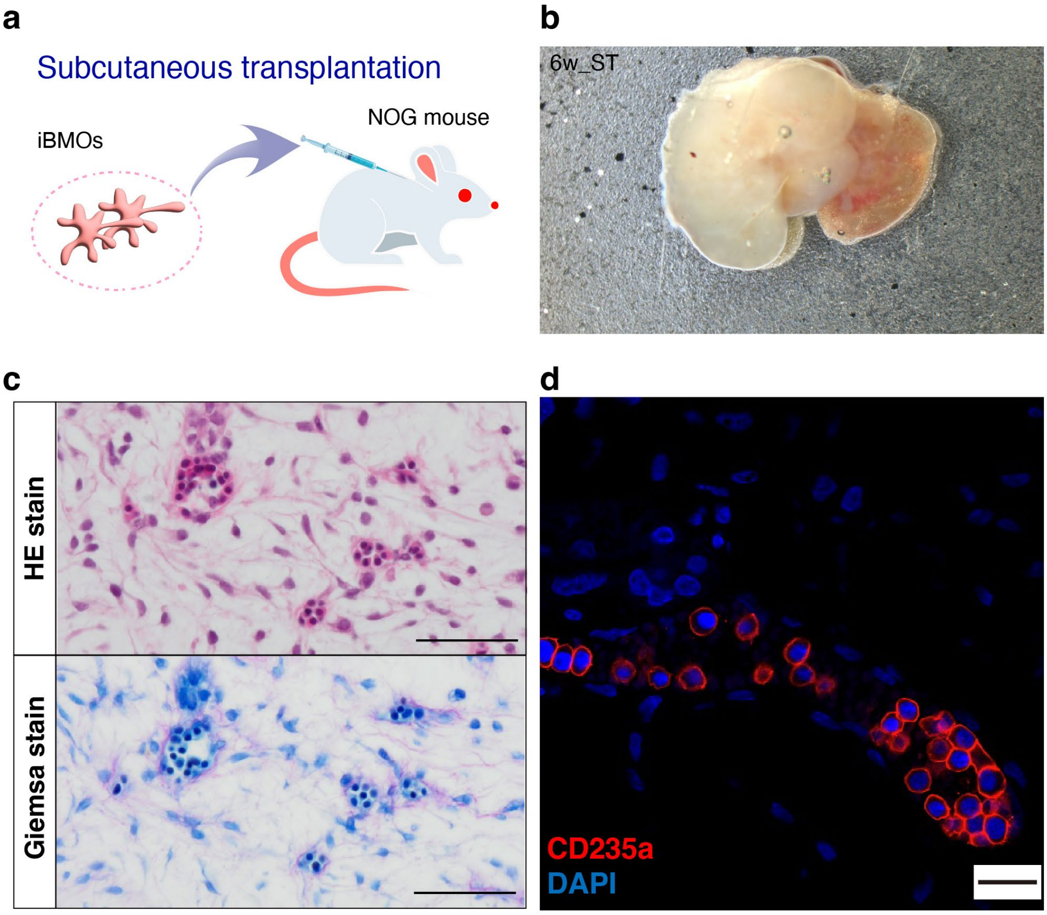


**Supplementary Figure 5. In vivo hematopoietic potential of human iBMOs.** **a.** Schematic of the transplantation strategy. Day 14 iBMOs were engrafted subcutaneously into immunodeficient NOG mice. **b.** Representative images of explanted grafts six weeks post-transplantation. **c.** Histology of subcutaneous (ST) grafts stained with hematoxylin and eosin (HE, top) and Giemsa (bottom). **d.** Immunofluorescence detected glycophorin A (CD235a) with DAPI nuclear counterstaining. Scale bars: 50 μm (C), 20 μm (D). ST, subcutaneous transplantation.
